# Neural effects of propofol-induced unconsciousness and its reversal using thalamic stimulation

**DOI:** 10.1101/2020.07.07.190132

**Authors:** André M. Bastos, Jacob A. Donoghue, Scott L. Brincat, Meredith Mahnke, Jorge Yanar, Josefina Correa, Ayan S. Waite, Mikael Lundqvist, Jefferson Roy, Emery N. Brown, Earl K. Miller

## Abstract

The specific circuit mechanisms through which anesthetics induce unconsciousness have not been completely characterized. We recorded neural activity from the frontal, parietal, and temporal cortices and thalamus while maintaining unconsciousness in non-human primates (NHPs) with the anesthetic propofol. Unconsciousness was marked by slow frequency (~1 Hz) oscillations in local field potentials, entrainment of local spiking to Up states alternating with Down states of little spiking, and decreased coherence in frequencies above 4 Hz. Thalamic stimulation “awakened” anesthetized NHPs and reversed the electrophysiologic features of unconsciousness. Unconsciousness is linked to cortical and thalamic slow frequency synchrony coupled with decreased spiking, and loss of higher-frequency dynamics. This may disrupt cortical communication/integration.

## Introduction

Propofol – the most widely used anesthetic - acts by enhancing GABAergic inhibition throughout the brain and central nervous system (Bai et al., 1999; Hapfelmeier et al., 2001; Hemmings et al., 2005). In humans, propofol produces dose-dependent changes in arousal level that are associated with spatiotemporal neurophysiological signatures across the cortex. At low doses, propofol produces a sedative state associated with beta oscillations (13-25 Hz) (McCarthy et al., 2008). At higher doses that maintain unconsciousness for surgery, slow-delta oscillation (0.1-4 Hz) (Lewis et al., 2012a; McCarley, 2007) appear across the entire scalp and are decoupled between cortical regions (Lewis et al., 2012a). Concomitantly, coherent alpha oscillations (8-12 Hz) concentrate across the frontal area of the scalp (Cimenser et al., 2011a; Purdon et al., 2013). In profound states of propofol-induced unconsciousness, the phase of the slow-delta oscillations strongly modulates the amplitude of the alpha oscillations (Purdon et al., 2013).

The broad range of dynamics observed in response to propofol administration suggests that propofol-induced unconsciousness is multifactorial process. One hypothesis is that unconsciousness is caused by the phase-locking of neuronal spiking with the slow-delta oscillation. This greatly reduces cortical spiking and limits cortical activity to brief Up-states of spiking followed by longer duration Down-states of little or no spiking (Lewis et al., 2012a). Another hypothesis is that the slow-delta oscillations “fragment” the cortex (Lewis et al., 2012a,b). Local spiking becomes limited to the narrow window of slow-delta oscillation phases, which being decoupled across cortical areas, impede long-range cortical communication. A third hypothesis, supported by modeling and experimental studies, suggests that the coherent frontal alpha oscillations represent hypersynchronous communication between the thalamus and prefrontal cortex (Ching et al., 2010; Palva and Palva, 2007). A fourth hypothesis links unconsciousness to loss of frontal-parietal connectivity (Lee et al., 2013). Finally, the neuroanatomy of the brainstem suggests that direct action of propofol at GABAergic inhibitory synapses onto the arousal center nuclei is also an important contributor to propofol-induced unconsciousness (Brown et al., 2010).

Details of propofol-induced unconsciousness remain to be clarified because its dynamics have been investigated mostly using extracranial measures such as electroencephalography (EEG) with limited spatial specificity and fMRI with limited temporal specificity (but see Redinbaugh et al., 2020). Microelectrode recordings in patients have highly precise temporal resolution but limited spatial coverage. Studies conducted simultaneously at high spatial and temporal resolution are needed. Moreover, most previous studies have not included the thalamus, a critical nexus that regulates cortical activity (Saalmann and Kastner, 2015; Sherman, 2016). The thalamus is highly interconnected with the cortex and receives important inputs from the brainstem arousal centers (Brown et al., 2010). Furthermore, both thalamocortical and corticothalamic connectivity are highly layer specific, giving rise to specific hypotheses that propose either deep (layers 5/6) or superficial (layer 2/3) may be more specifically linked to loss of consciousness (Aru et al., 2019; Dehaene and Changeux, 2011). Dehaene and Changeux (2011) link this to frontal cortex. By contrast, others link consciousness to parietal and posterior cortex (Boly et al., 2017). Central thalamic stimulation can result in a partial restoration of consciousness (Schiff et al., 2007a). Some theories suggest consciousness is formed when oscillations in thalamocortical loops integrate cortical information (Llinás, 2003). There is mounting evidence that the thalamus regulates cortical communication (especially top-down) via oscillatory dynamics (Halassa and Sherman, 2019; Saalmann and Kastner, 2015).

To study the effects of propofol on cortex and thalamus, we implanted vascular access ports subcutaneously in four non-human primates (NHPs). This allowed us to administer propofol using computer-controlled infusion and continuously record neural activity as the animals transitioned from the awake state, through loss of consciousness (LOC), to unconsciousness and recovery of consciousness (ROC) (Fig, 1A).

**Figure 1.**
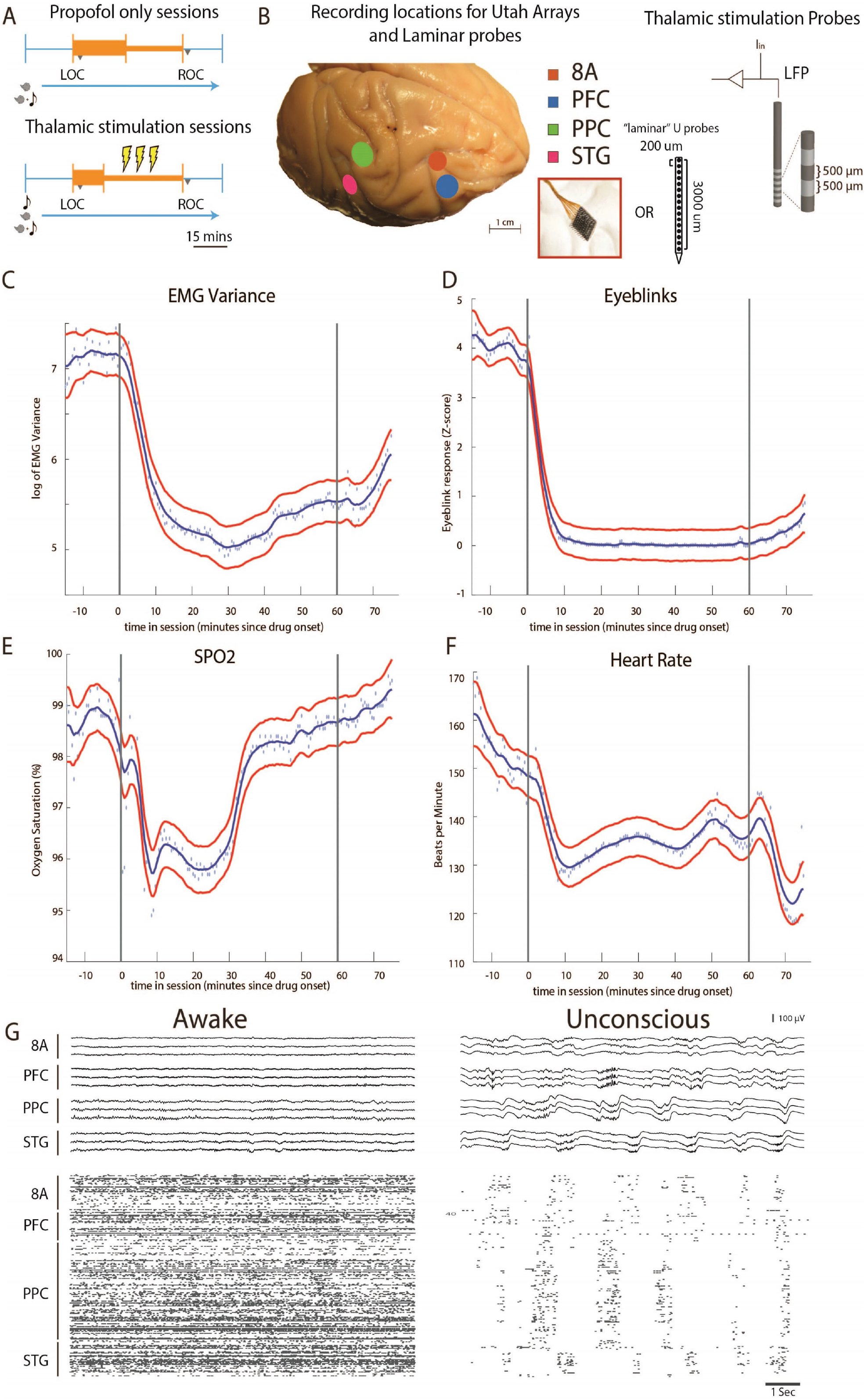
Propofol anesthesia paradigm and physiological indices of LOC. A, Session paradigm. Two sets of sessions were performed, for propofol only sessions (upper subplot), there was an initial 30-minute infusion (fast rate, thick orange bar) covering awake (pre-LOC) and generally anesthetized epochs, before switching to a halved rate propofol infusion for maintenance phase of experiment (narrow orange bar). For the thalamic stimulation sessions (N=22), the initial infusion was for 20 minutes, followed by a halved rate propofol infusion for the rest of the session. Periodically, 30-second DBS trials (yellow bolts) occur during lower-dosed maintenance phase of propofol infusion. LOC: Loss of consciousness, ROC: Recovery of consciousness. B, (left) Cortical recording locations of each 64 channel chronic recording arrays or 16 channel acutely inserted laminar probes. PFC: ventrolateral prefrontal cortex; 8A: caudal lateral PFC; PPC: posterior parietal cortex; STG: superior temporal gyrus; (right) Thalamic electrical stimulation leads. C-F, Physiological measurements characterizing the pre-drug awake state relative to Propofol administration (starting at time zero). Dots indicate individual time points, with measurements averaged over sessions, red lines indicate the 95% Confidence Interval (see Methods), and blue lines are a smoothed version of the data. G, (upper panel) Example LFP traces from all cortical arrays during the awake (left) and post-LOC states with clear Slow Frequency waves (right). (lower panel) Example spike raster over 10 seconds of data. Spike times are indicated with dots.

## Results

### Experimental design, recordings, and physiological responses to propofol

We started by administering a high-infusion rate of propofol (280-580 mcg/kg/min, adjusted per individual animal) for 20 minutes to induce LOC (defined as the moment the eyes closed). They remained closed for the remainder of the infusion. LOC was associated with simultaneous changes in a several physiological variables. We assessed the significance of the change in each physiological variable as the posterior probability that its values were greater during the baseline state (prior to initiation the propofol infusion) than they were 10 to 60 minutes after initiation of the propofol infusion (Smith et al. 2010). We considered a change to be significant if the posterior probability was greater than 0.99.

Relative to the pre-propofol awake state, we observed a significant decrease in muscle tone (from 4.5 to 75 minutes, Figure 1C), cessation of airpuff-evoked eyeblinks (from 1 to 75 minutes, Figure 1D), a decline in O_2_ saturation (from 4.5 to 34 minutes, Figure 1E), and a decrease in heart rate (from 4.5 to 75 minutes, Figure 1F). All times were measured relative to the start of the propofol infusion. After LOC, we decreased the infusion to a maintenance rate (140-230 mcg/kg/min) for an additional 30-45 minutes. Once the infusion was ceased, return of consciousness (ROC) occurred in approximately ~7 minutes. ROC was behaviorally defined as the moment the eyes opened and remained open continuously. We defined unconsciousness as the period between LOC and ROC.

We performed the neurophysiological recordings in three sets of experiments. In the first set of experiments, we recorded neuronal spiking activity and local field potentials (LFPs) from a series of chronically implanted 64-channel “Utah” arrays in frontal cortex (8A, PFC), posterior parietal (PPC, 7A/B), and auditory/temporal (superior temporal gyrus, STG) cortex. These initial sessions (ten in monkey 1, 11 in monkey 2) served to establish the neurophysiological properties defining the awake state and the state of unconsciousness. In the second set of experiments, we implanted multiple-contact stimulating electrodes (two in each hemisphere of the same monkeys) in frontal thalamic nuclei (intralaminar nuclei, ILN, and mediodorsal nuclei, MD) to record from and electrically stimulate the thalamus and cortex during propofol-induced unconsciousness (a total of 22 additional sessions, 11 in monkey 1, 11 in monkey 2). In the third set of experiments, in two additional monkeys (a total of eight sessions, two in monkey 3 and six in monkey 4), we performed acute laminar recordings from the same areas using multi-contact arrays positioned approximately perpendicular to cortex (as in Bastos et al., 2018).

An example session of cortical recordings during the pre-propofol awake state, and unconsciousness (Figure 1G) showed a profound reduction in spiking, an increase in both slow-frequency LFP amplitude and slow-frequency modulation of spiking activity.

### Changes in LFP power with unconsciousness

We first characterized the changes in LFP power during the pre-propofol awake state (− 20 to −15 minutes pre-propofol infusion) to every time point after drug administration until 10 minutes post-ROC (non-parametric cluster-based randomizations, corrected for multiple comparisons; all effects p<0.01, (Maris and Oostenveld, 2007)). We compared LFP power increases/decreases from the awake state, time locked either to LOC or ROC. First, approximately 3-7 minutes before LOC (alternatively, 3-7 minutes after beginning propofol infusion), there was an increase in beta power in all areas (~15-30 Hz). In frontal areas PFC and 8A, this power change shifted to a lower frequency (~14-20 Hz) during the maintenance phase (Figure 2A and B, left subplots). Power in this frequency range remained higher than awake throughout unconsciousness. By contrast, in posterior areas PPC and STG after LOC (dotted black lines at time point 0 in Figure 2, left subplots) beta power was reduced relative to awake (Figure 2C and D, left subplots). In all areas, shortly before LOC there was a sustained reduction in gamma power (>35 Hz) and an increase in slow-frequency (SF, 0.1-3 Hz) power.

**Figure 2.**
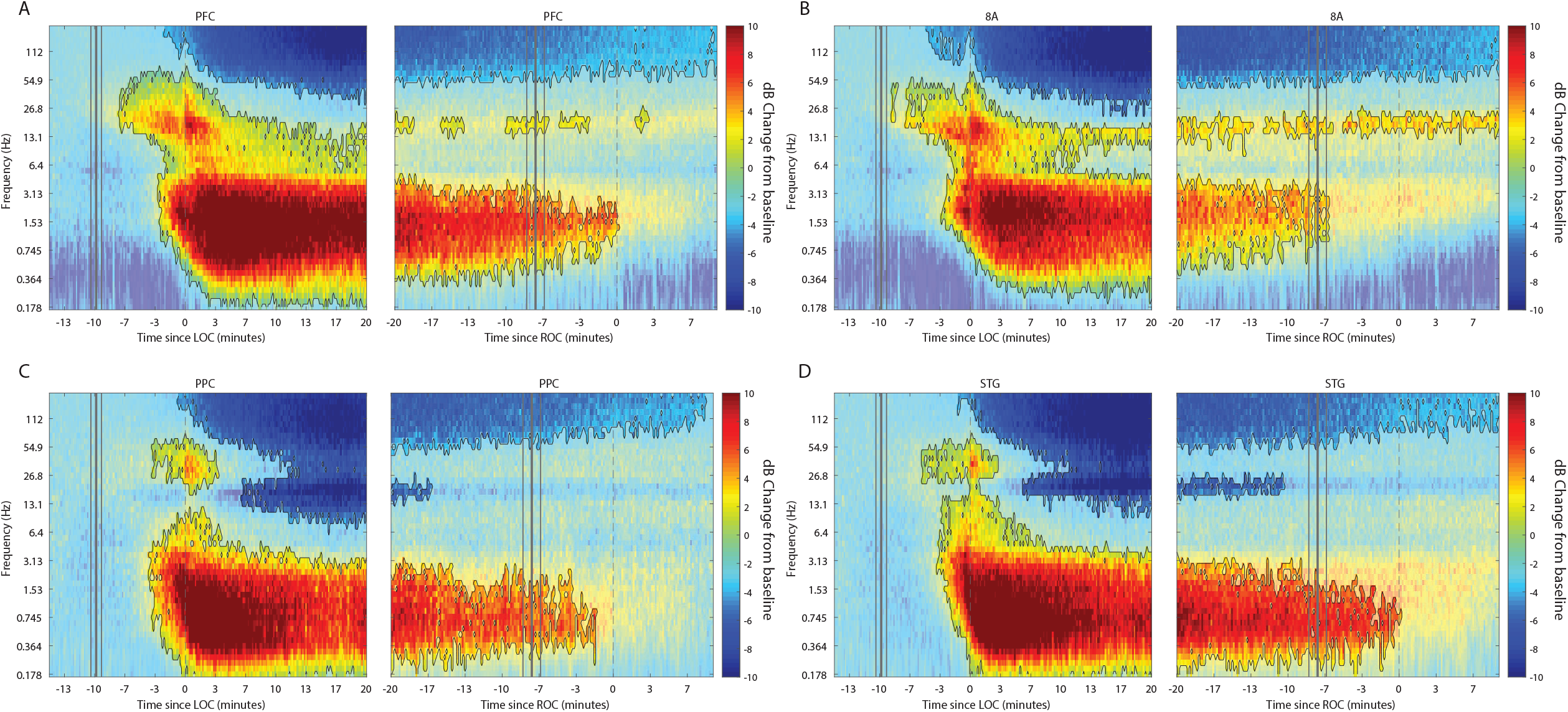
Changes in power during unconsciousness. A-D dB change in power for each area is shown with respect to pre-propofol awake baseline. Increases in power during unconsciousness are shown in red, decreases in blue. Significant modulation of power is shown in opaque colors and outlined in black. Left subpanels are time locked from Loss of Consciousness (LOC), defined behaviorally as the moment the eyes closed and remained closed. Start of drug infusion is shown as a vertical black bar at −10 minutes from LOC. Right subpanels are time locked from Return of Consciousness (ROC), defined behaviorally as the moment the eyes first opened. Cessation of drug infusion is shown as a vertical black bar at −8 minutes from ROC.

We quantified the effect size by comparing power while awake to power during unconsciousness (Supplemental Figure 1A, blue and orange bars, respectively). We measured the effect size in dB units (10*log_10_(power awake/power unconscious) across areas. In the slow frequency range, the order of areas from strongest to weakest change was STG, PPC, PFC, and 8A (Supplemental Figure 1A/C, lower subpanel, significant differences in effects sizes were observed between STG and PFC and STG and 8A, P<0.01, non-parametric randomization test).

### Anteriorization of alpha/beta power and differences in slow-frequencies across areas during unconsciousness

“Alpha anteriorization” is the decrease in alpha/beta power in posterior cortex, and its increase in frontal cortex, during unconsciousness (Cimenser et al., 2011b; Jh et al., 1977). We plotted the effect size (10*log_10_(power awake/power unconscious)) as a function of frequency (Figure 3A). This revealed differences between posterior and anterior areas in the alpha/low beta range. In particular, area 8A showed a clear peak at 15 Hz during unconsciousness (Figure 3A, red line). At that frequency and above, posterior brain regions were strongly depressed in power (Figure 3A, green and magenta lines). We directly compared the absolute power in all areas between the awake and the unconscious states. We found that in the awake state, alpha/beta (15-25 Hz) power in posterior areas PPC and STG was greater than in anterior areas (Figure 3B, sign test for anterior vs. posterior alpha/beta power across sessions, p<1E-3 for all comparisons). During unconsciousness this relationship flipped. Alpha/beta power in frontal areas PFC and 8A was higher than posterior areas (Figure 3C, sign test for anterior vs. posterior alpha/beta power across sessions, p<1E-5 for all comparisons).

**Figure 3.**
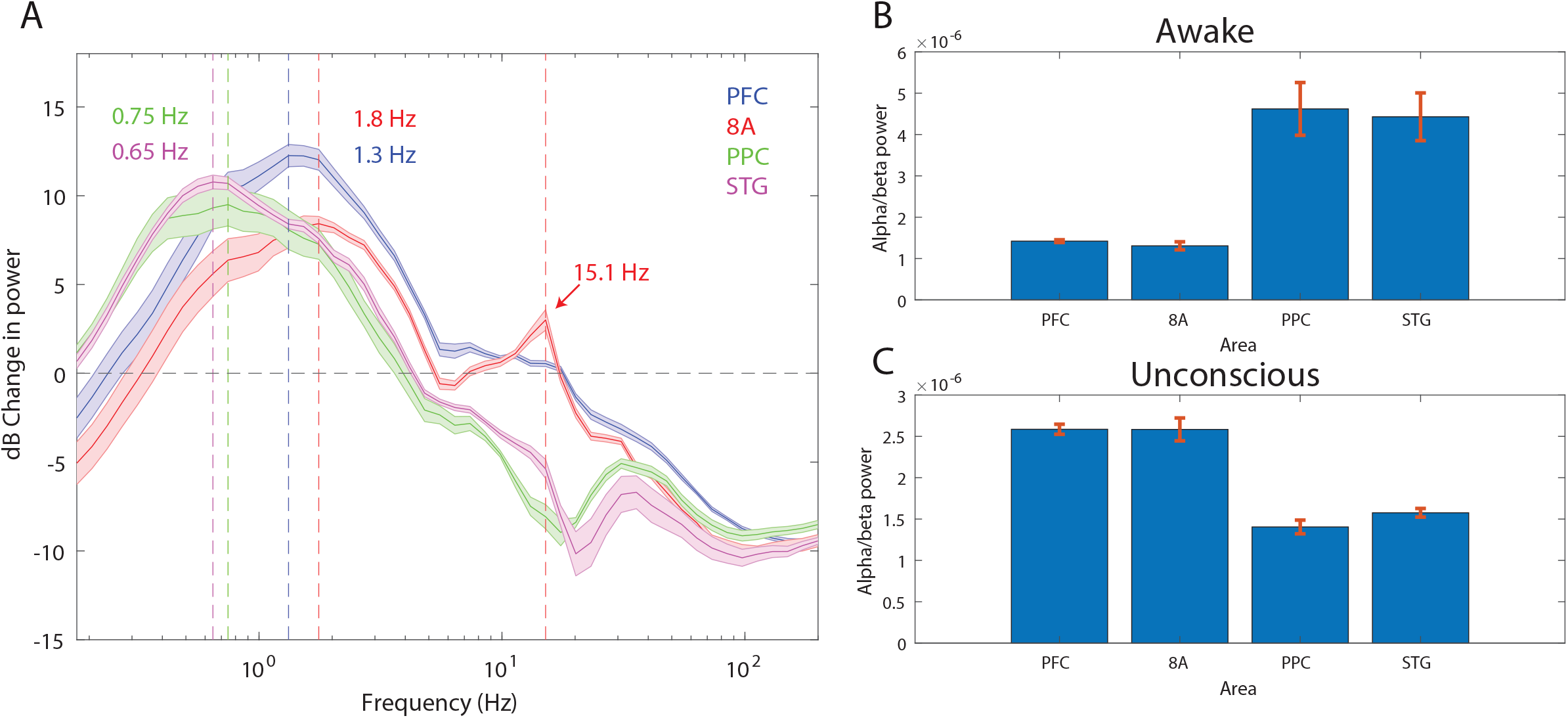
Power asymmetries between areas. A, dB Change in power during all time points of unconsciousness vs. pre-propofol awake baseline. For each area, the mean power difference is shown +/− 1 SEM. The Slow Frequency peaks are highlighted for each area, and a secondary peak in the beta range (15.1 Hz) is present in area 8A. B, Total power in the beta frequency range (15-25 Hz) during pre-propofol awake baseline. C, Same as B, but during unconsciousness. Mean +/− 1SEM.

This analysis revealed differences in which frequencies of the slow frequency band changed the most between awake and unconscious. In anterior areas, PFC and 8A, the peak change was in higher frequencies (Figure 3A, 1.3 and 1.8 Hz, respectively) than posterior areas, PPC and STG (Figure 3A, 0.75 and 0.65 Hz, respectively, sign test for frequency difference, all anterior vs. posterior comparisons, p<1E-5).

### LFP power changes from awake to unconscious

After cessation of propofol infusion, ROC occurred ~8 minutes later. In all areas, at ROC, power in the slow frequency range was no longer significantly different from the awake state (Figure 2A-D, right subpanels). In contrast, significant reductions in gamma power were still present in PFC, 8A, and STG even 10 minutes post-ROC. Beta power changes from awake were also not reliable indicators of ROC. In frontal area 8A, beta power remained elevated above the awake-state levels 10 minutes post-ROC (Figure 2B, right subpanel). In posterior areas, beta power reductions from awake were no longer significant at 10-17 minutes pre-ROC.

### Differences in spiking between awake vs. unconscious

We next examined differences in spike rates between the awake and unconscious states (Figure 4A). We eliminated periods with airpuffs or auditory tones in order to not induce differences across areas due to differences in their sensory processing. In the awake state, average spike rates were between 6-8 spikes per second across areas. During the unconscious state, average spike rates across areas fell to 0.2-0.5 spike/sec during the initial infusion (first 30 minutes of propofol). They increased to 1-2 spikes/sec during the maintenance infusion. After cessation of propofol, spike rates gradually approached the levels seen in the awake state (Figure 4A).

**Figure 4.**
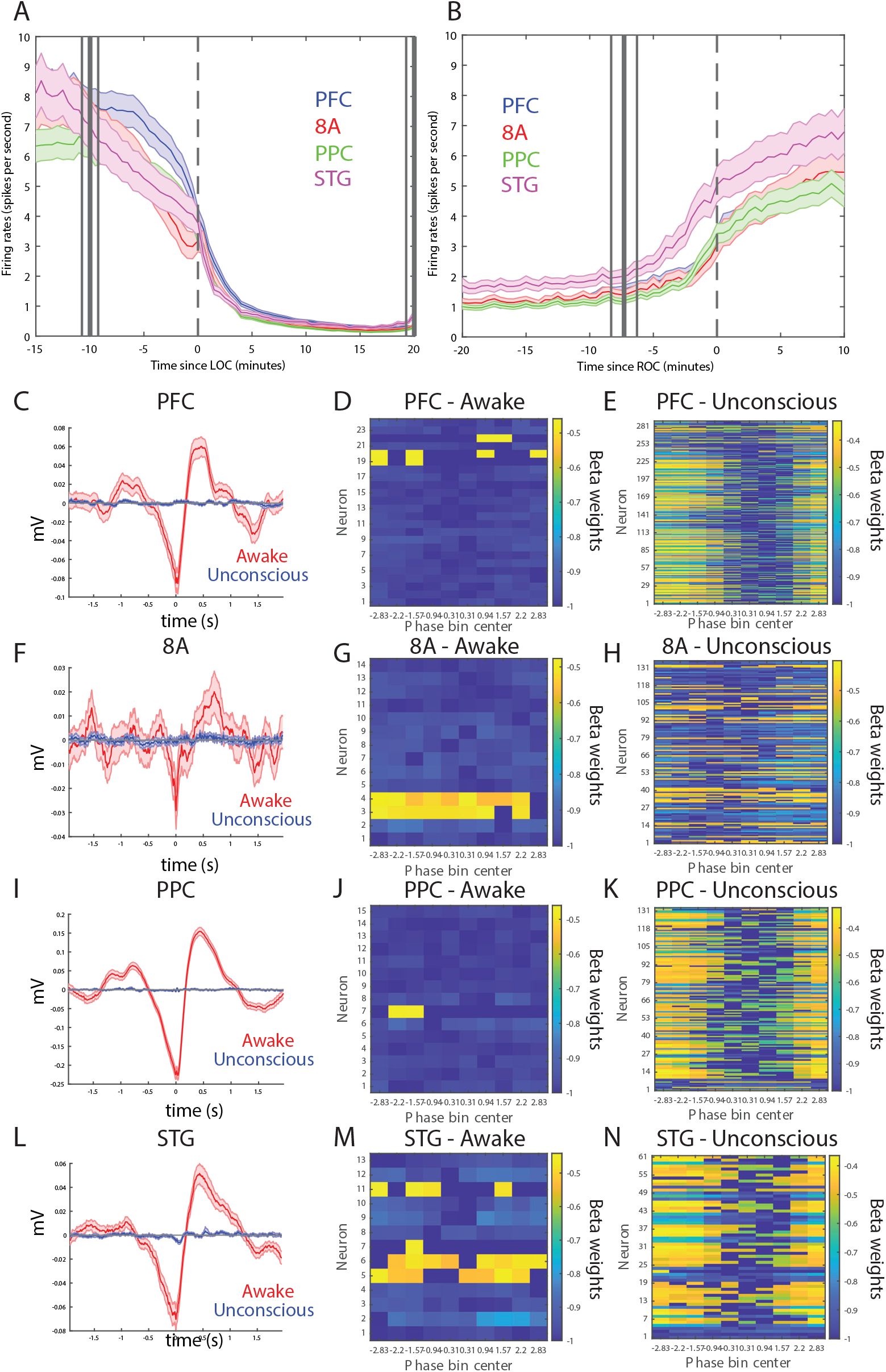
Changes in mean firing rate and spike-field coupling during awake vs. unconscious states. A, Spike rate for all recorded areas averaged across all Propofol-only recording sessions locked to LOC. B, Same as A, but for ROC. Mean and 99^th^ confidence interval. C/F/I/L The spike triggered average for all well-isolated units in a given area with respect to that area’s unfiltered LFP. Red is the awake state, blue is the unconscious state. D/G/J/M Beta weights for all units with significant coupling to the SF (0.1-3 Hz) LFP during the awake state. E/H/K/N Beta weights for all units with significant coupling to the SF (0.1-3 Hz) LFP during the unconscious state.

At LOC, spike rates were approximately half of their awake rates (Figure 4A, mean firing rates at LOC: 4.0, 3.2, 3.4, 3.8 spikes/sec for PFC, 8A, PPC, STG, respectively). Spike rates continued to fall and then stabilized ~15 minutes post-LOC to an average of 0.25 spikes/second. During the maintenance infusion, average spike rates across areas increased to 1-1.68 spikes/second.

At ROC, spiking activity increased to 3.3, 3.0, 3.4, and 5.0 spikes per second (for PFC, 8A, PPC, STG, respectively), approximately the same spike rate as at LOC (Figure 4B). The exception was area STG, which recovered faster than the other areas. Neurons in STG had an average of 5.0 spikes per second at ROC, significantly greater than the other areas (p<0.01, non-parametric randomization test for all area comparisons to STG).

We quantified the effect size by comparing spike rates during the awake state to the unconscious state (Supplemental Figure 1B). We measured the effect size in dB units (10.*log_10_(firing rate awake/spike rate unconscious) across areas. The order of areas from highest to lowest spike rate change was hierarchically organized. The strongest effects were in PFC/8A, then PPC, then STG (Supplemental Figure 1D, lower subpanel, all comparisons, p<0.01, non-parametric randomization test).

During unconsciousness, spike timing was phase-coupled to the slow frequencies. This appeared as Up and Down states of high vs minimal/no spiking (e.g., Figure 1G) respectively. Spike-triggered averages of the local LFP signal indicated that spikes entrained to the depolarized phases (troughs) of slow frequency oscillations (Figure 4C/F/I/L) in all areas. We further tested whether spikes could be predicted as a linear combination of the LFP slow-frequency/delta oscillation phase using a generalized linear model (GLM) framework. Figure 4D/G/J/M (4E/H/K/N) shows the model coefficients for each neuron in each area during the awake (unconscious) state. Only coefficients corresponding to models with a good fit to the data are shown (see Methods). Using this framework, we consider a neuron to be phase-modulated if this model describes well the data. We compared the proportion of phase-modulated neurons during the unconscious state to the proportion of phase-modulated neurons during the awake state by performing a Bayesian comparison, using the beta-binomial model (DeGroot and Schervish, 2012). The posterior probability that the proportion of phase-modulated neurons was greater during the unconscious state than during the awake state was greater than 0.99 for all brain regions.

### Laminar changes in spiking and LFP power during unconsciousness

In two additional animals we used multi-laminar probes to examine differences between cell layers (Figure 5). We pooled data from PFC, 8A, PPC, and STG because these frequencies behaved consistently across areas. Laminar position zero was based on the relative power profile of gamma vs. beta (see Methods). This was previously shown to map onto the location of layer 4 (Bastos et al., 2018). As before, we calculated the dB change from the awake state to unconsciousness. The laminar profile of change in spiking, gamma, and slow frequencies are shown in Figure 5A/C/E. Spiking and gamma reduction were more pronounced in superficial layers compared to deep layers (Figure 5B and D, non-parametric randomization test comparing the effect size of superficial vs deep, P<1E-7 for spiking, P<1E-7 for gamma, (Manly, 2018)). The enhancement of slow frequency power was higher in deep compared to superficial layers (Figure 5F, non-parametric randomization test comparing the effect size of superficial vs deep, P<1E-5 for SF).

**Figure 5.**
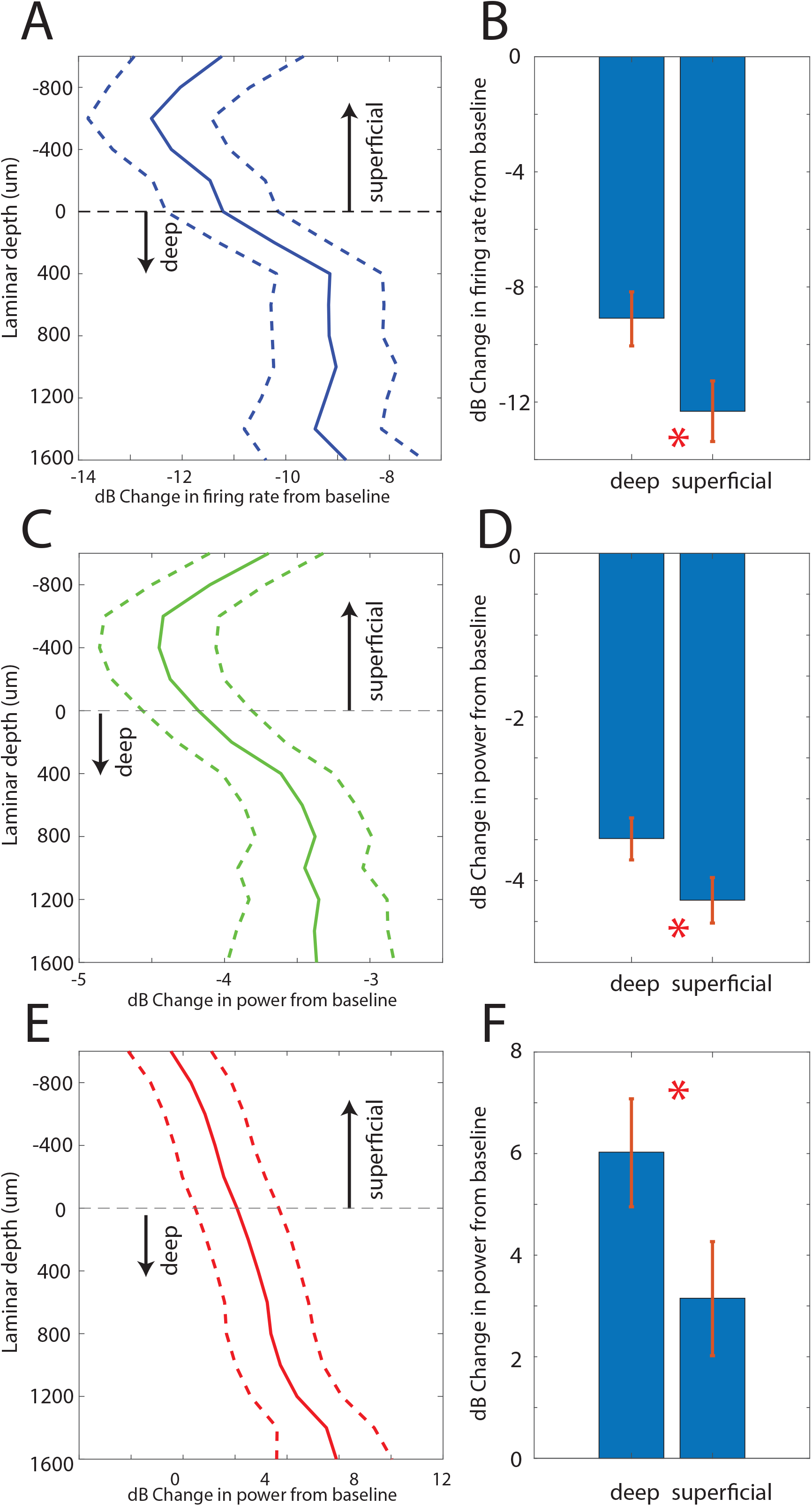
Laminar changes in spiking, gamma, and SF power during unconsciousness. A, Firing rate change from pre-propofol baseline as a function of cortical layer. Layer 0 is the approximate location of cortical layer 4 (see Methods). The horizontal dotted line at zero separates superficial layers 2/3 from deep layers 5/6. The average effect size in firing rates per layer comparing unconsciousness (LOC-30 minutes post-propofol onset) to pre-propofol baseline. B, Mean and SEM of the effect size across all superficial (N=287) and deep (N=337) neurons. C, D, same as A, B, but for LFP power at Gamma (100-200 Hz) E, F, Same as A, B, but for SF power (0.2-1.1 Hz), respectively (N = 330 LFPs for superficial layers, N = 393 LFPs for deep layers). Mean and 99^th^ Confidence Interval across all available units in each layer

### Changes in corticocortical and thalamocortical coherence during unconsciousness

We next analyzed whether slow frequency LFP oscillations were also phase synchronized. We used a sliding window approach to quantify the Pairwise Phase Consistency (Vinck et al., 2010). Pairwise Phase Consistency is a metric of phase synchronization that is un-biased by the number of observations. There were strong increases in slow frequency cortico-cortical phase synchronization between all cortical areas (P<0.01, cluster-based non-parametric randomization test, Figure 6) during unconsciousness. In a subset of areas (Figure 6B/E/F), there were reductions in cortico-cortical theta and beta phase synchronization, although the effect size at these frequencies was smaller than at slow frequencies (Supplemental Figure 2). Unique to phase synchronization between frontal areas PFC and 8A, we observed a sustained increase in alpha (7-15 Hz) frequency range phase synchronization during unconsciousness (P<0.01, cluster-based non-parametric randomization test, Figure 6A).

**Figure 6.**
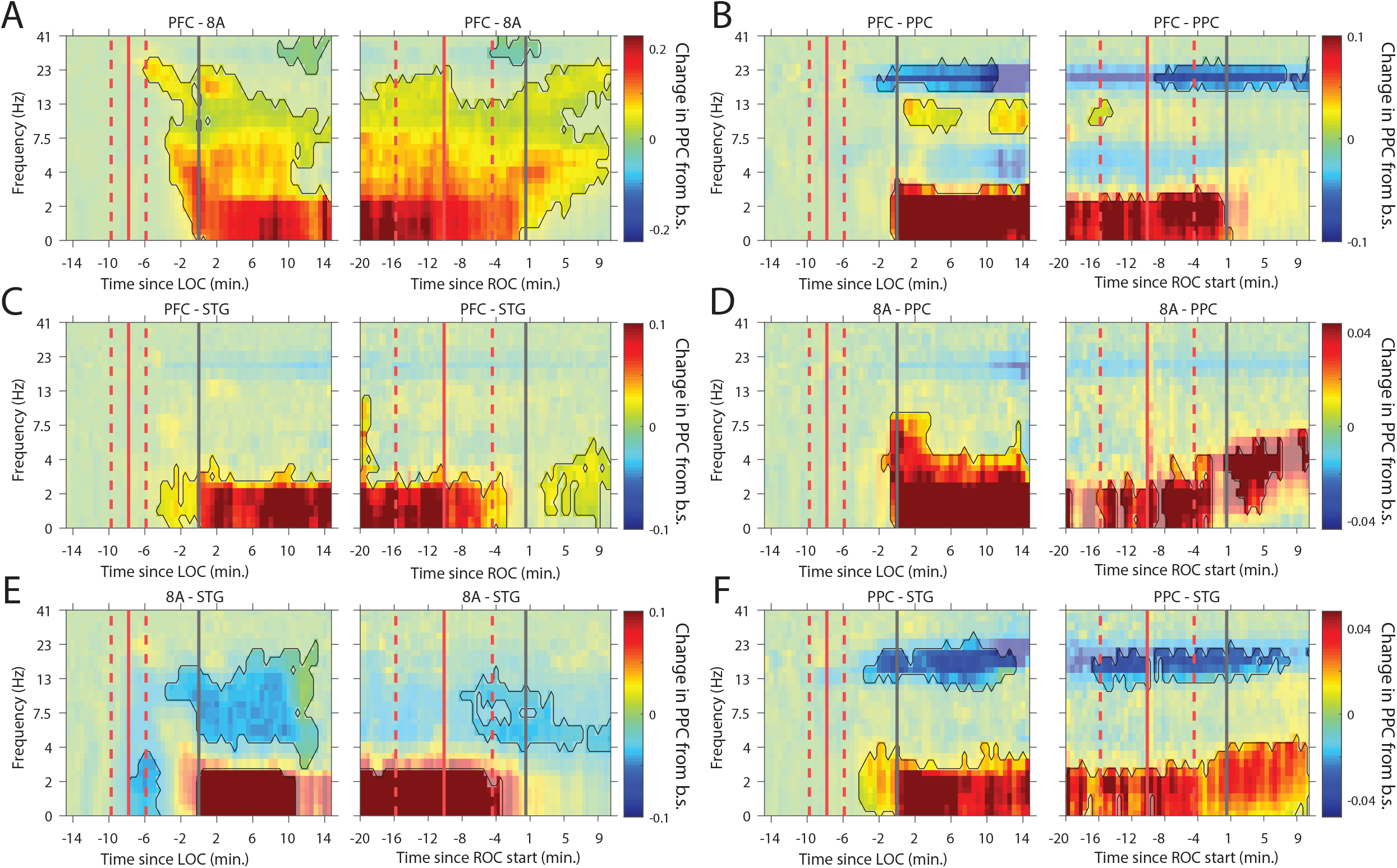
Changes in cortico-cortical coherence during awake vs. unconscious states. A-F (left subpanels) Change in the Pairwise Phase Consistency (PPC) for all time points relative to LOC compared to pre-propofol awake baseline (−15 to −10 minute pre-LOC). The red vertical lines indicate the average +/− 1 standard deviation time of propofol onset. The black vertical line indicates time zero (LOC). Significant increases or decreases from awake baseline are opaque colors and are highlighted. (right subpanels) Same as left subpanels but for ROC. The red vertical lines indicate the average +/− 1 standard deviation time of propofol offset. The black vertical line indicates time zero (ROC). Significant increases or decreases from awake baseline are opaque colors and are highlighted.

Recent evidence suggests that the thalamus helps foster synchrony between cortical areas (Saalman and Kastner, 2015). During unconsciousness, all cortical areas show increased slow frequency phase synchronization with the thalamus (P<0.01, cluster-based non-parametric randomization test, Figure 7). There were also reductions in synchrony during unconsciousness in the theta/beta frequency ranges, but as with cortico-cortical phase synchronization, these effects were smaller (Supplementral Figure 3). Uniquely to phase synchronization between thalamus and PFC, we observed a sustained increase in alpha (7-15 Hz) frequency range phase synchrony during unconsciousness (P<0.01, cluster-based non-parametric randomization test, Figure 7A).

**Figure 7.**
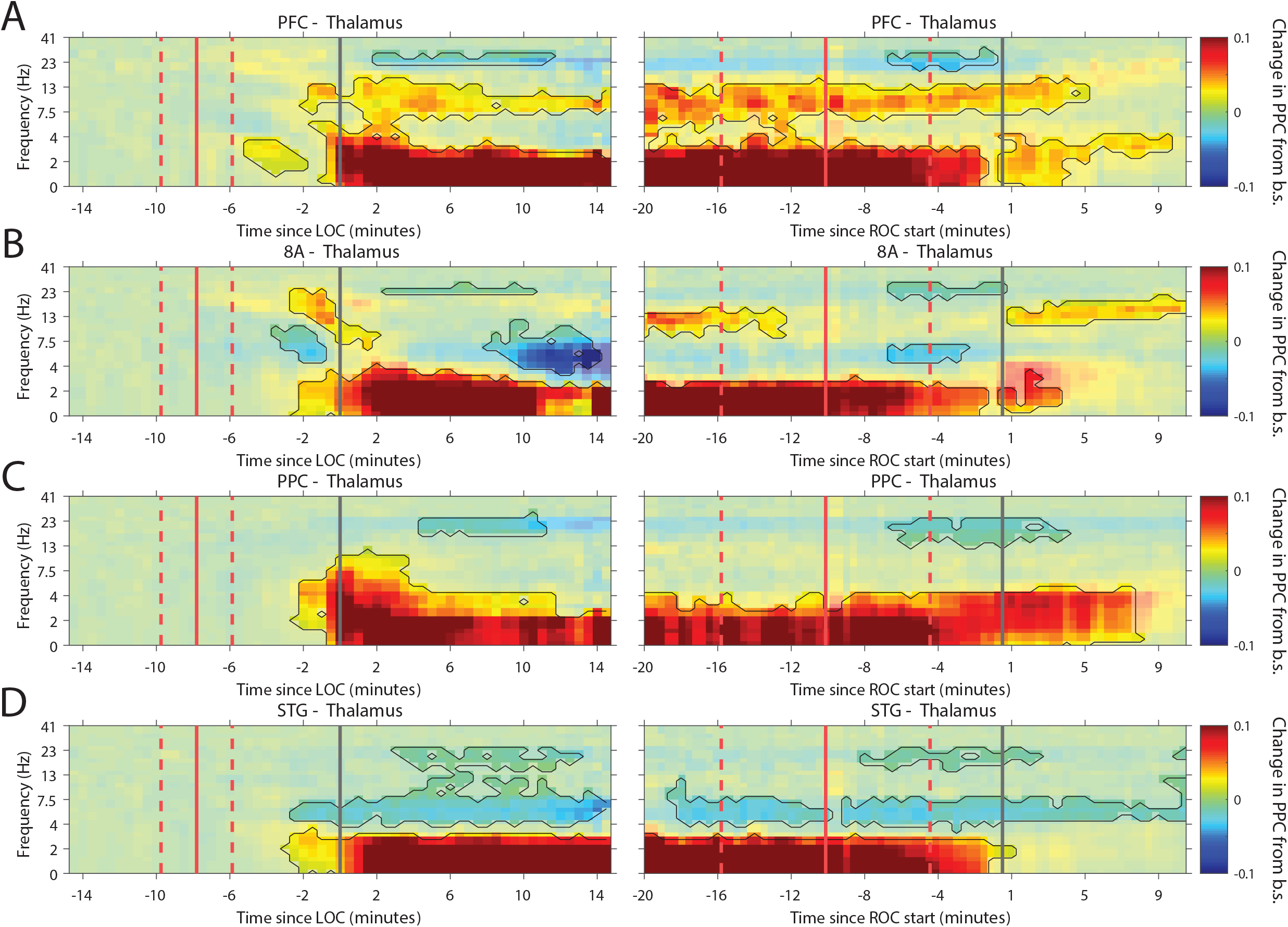
Changes in thalamo-cortical coherence during awake vs. unconscious states. A-D (left subpanels) Change in the Pairwise Phase Consistency (PPC) for all time points relative to LOC compared to pre-propofol awake baseline (−15 to −10 minute pre-LOC). The red vertical lines indicate the average +/− 1 standard deviation time of propofol onset. The black vertical line indicates time zero (LOC). Significant increases or decreases from awake baseline are marked with opaque colors and are highlighted. (right subpanels) Same as left subpanels but for ROC. The red vertical lines indicate the average +/− 1 standard deviation time of propofol offset. The black vertical line indicates time zero (ROC). Significant increases or decreases from awake baseline are opaque colors and are highlighted.

### Thalamic stimulation arouses unconscious monkeys and causes a partial reversal of neurophysiological signs of unconsciousness

The thalamus is a major route by which ascending excitatory projections from the brainstem reach the cortex. We tested whether we could induce behavioral arousal and restore awake-like neurophysiological markers by electrically stimulating the thalamus. During unconsciousness, we applied 180 Hz, bipolar stimulation targeting the central thalamus, including the mediodorsal nucleus and intralaminar nuclei (Figure 8B, see Methods). We applied either high or low current, titrating it for each individual animal (Figure 8A) based on the change in arousal that it evoked. We measured arousal with a “wakeup score” (see Methods). It assessed whether the eyes opened and whether there was an increase in limb movements and puff-evoked eyeblinks (see Methods). A wakeup score of zero indicated none of these happened. A value of 6 indicated that they all occurred. During unconsciousness without electrical stimulation, we did not observe these events (e.g., Figure 1D shows that on average there were no air puff evoked eyeblinks during unconsciousness). Analysis of the EMG, SPO2, eyeblinks to airpuffs, and heart rate during thalamic electrical stimulation corroborated this wakeup score: Electrical stimulation of the thalamus increased muscle tone (non-parametric randomization test, p<0.01, Figure 8C), eyeblinks to airpuffs (non-parametric randomization test, p<0.01, Figure 8D), blood oxygenation (non-parametric randomization test, p<0.01, Figure 8E), and heart rate (non-parametric randomization test, p<0.01, Figure 8F). These changes were all greater for high relative to low current (Figure 8C-F, non-parametric randomization test for significant differences between high vs. low current stimulation, P<0.01 are indicated with black stars) and outlasted the electrical stimulation period itself, achieving significance during the time window from 0-30 seconds post stimulation offset (post-stim 1 in Figure 8C-F) and sometimes the 30-60 seconds (post-stim 2) or 60-90 seconds (post-stim 3) post stimulation offset periods.

**Figure 8.**
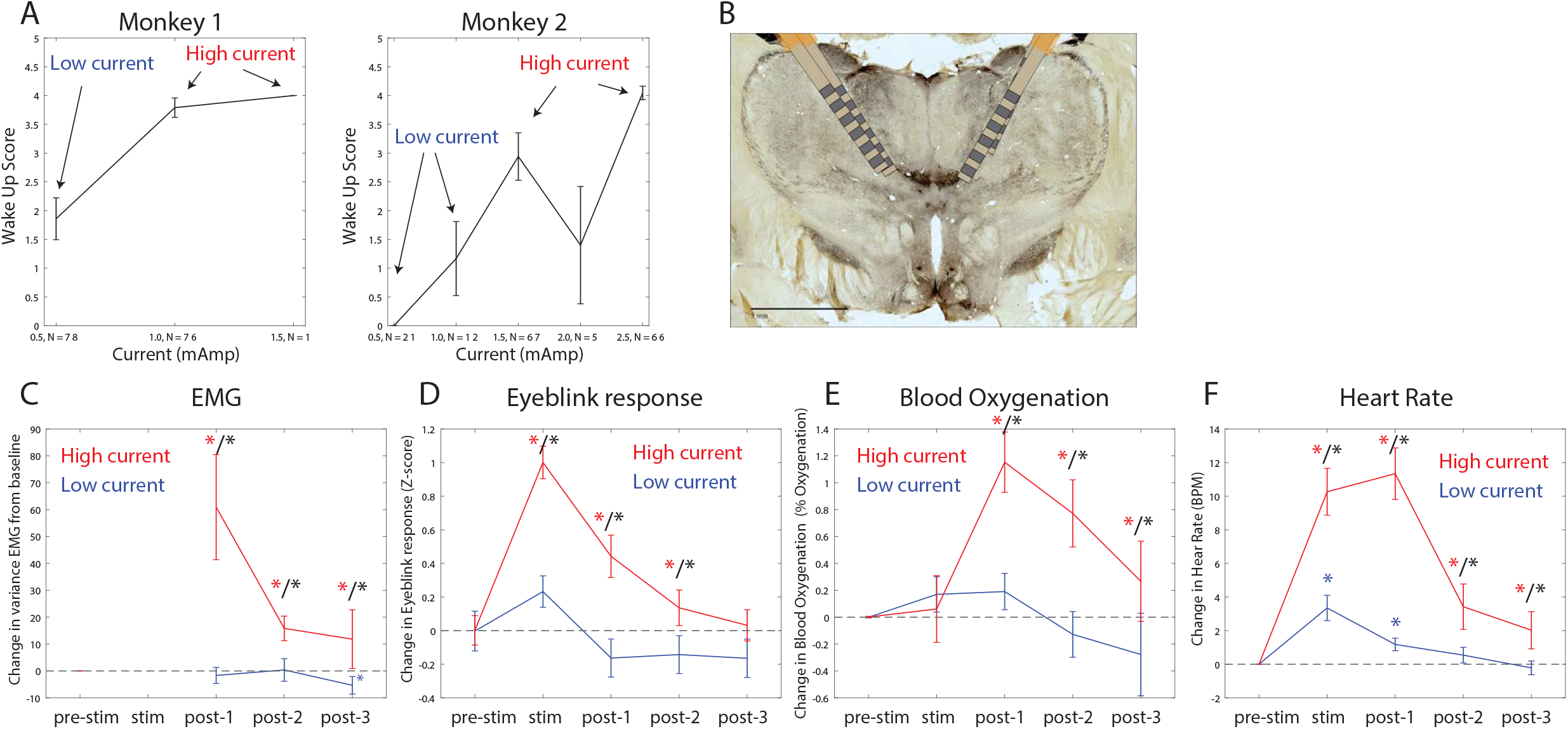
Thalamic electrical stimulation in central thalamus arouses monkeys. A, Behavioral wake-up score as a function of thalamic current for monkey 1. Mean +/− 1 SEM. A high current and low current condition was individually titrated per monkey for producing scores on average above or below a wake-up score of 2. B, A histological image from monkey 1 showing the thalamic leads in the central thalamus. C, Change in EMG from pre-stimulation baseline for high current (red) vs. low current (blue) conditions. Change in the physiological signal was tested for difference from zero during the stimulation period (0-28.5 seconds with respect to electrical stimulation onset), post-1 (0-30 seconds with respect to electrical stimulation offset), post-2 (30-60 seconds with respect to electrical stimulation offset), and post-3 (60-90 seconds with respect to electrical stimulation offset). Significant differences from zero are indicated with red or blue stars. Significant differences in high vs. low current are indicated with black stars. Mean +/− 1 SEM. D, same as C, but for Eyeblink response to air puffs. E, Same as C, but for Blood Oxygenation. F, same as C but for the heart rate response.

Stimulation produced an awake-like cortical state by increasing spiking rates and decreasing slow frequency power. An example raster plot of well-isolated single neurons from a single stimulation trial is shown in Figure 9A. To see how we removed spurious threshold crossings due to the electrical stimulation itself, see Methods. High-current stimulation increased the spike rate (Figure 9B and Figure 9C, upper subpanel) from ~1 to ~2.5 spikes per second. In all areas, there was a significant increase in spiking during the stimulation interval compared to pre-stimulation baseline (Figure 9C, orange bars in upper subpanel, red stars indicate significant differences, P<0.01, non-parametric randomization test). In area STG, this increase brought the average spike rate to the same level as that seen during natural ROC (horizontal dotted lines in Figure 9C, upper subpanel). In all areas, the increased spike rate persisted for at least 30 seconds (and as long as 90 seconds) post-stimulation (Figure 9C, red stars indicate significant differences, P<0.01, non-parametric randomization test). With low current stimulation, mean spikes rates were unchanged from pre-stimulation baseline (Figure 9C, lower subpanel, all P>0.05).

**Figure 9.**
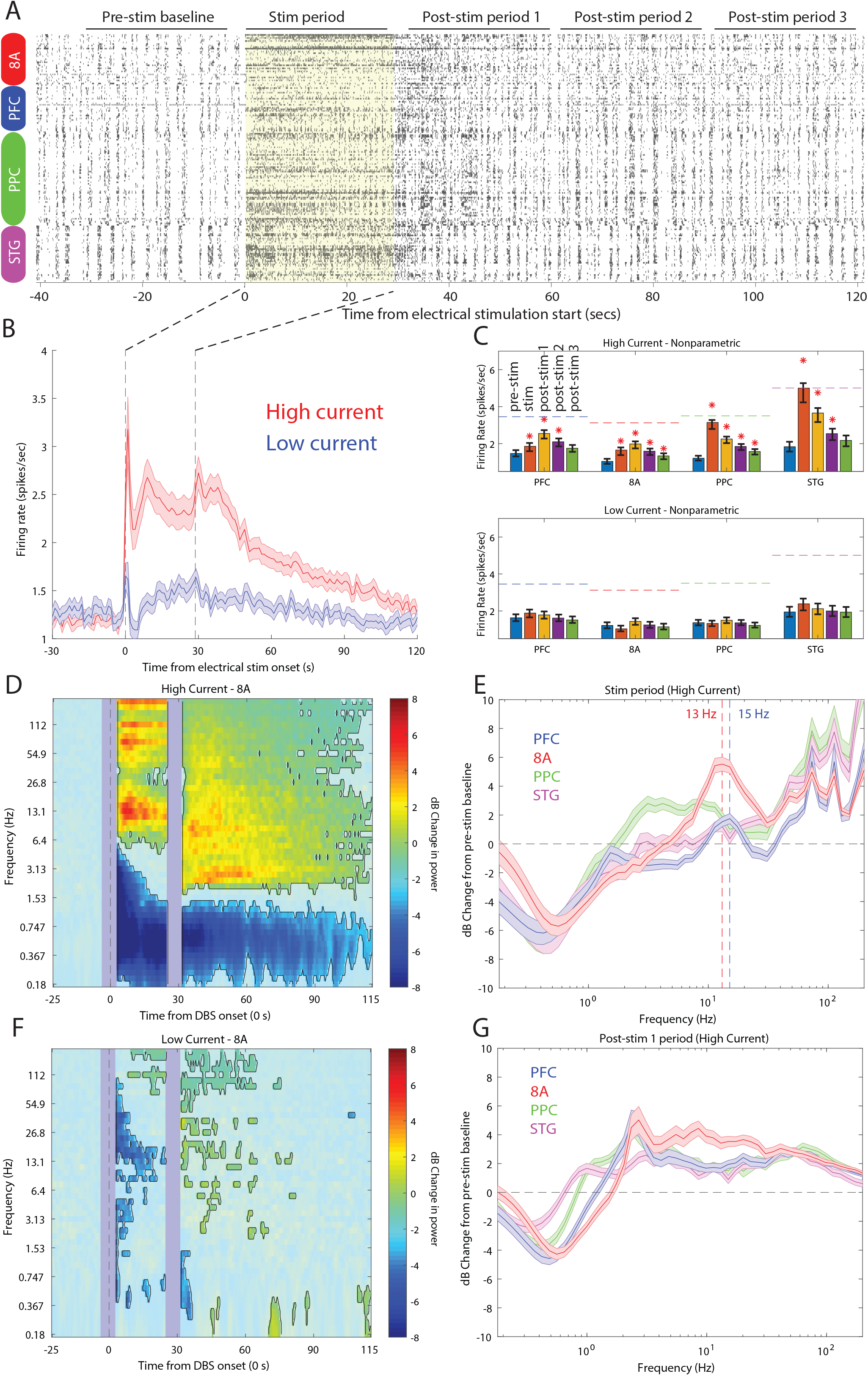
Effects of thalamic electrical stimulation on cortical state. A, An example trial, well-isolated single units are shown before, during, and after electrical stimulation for a trial that produced a maximal wake-up score (eyes opened, muscle movement, response to air puffs). B, The average effect on all well-isolated single units across all areas. Mean firing rates with respect to electrical stimulation onset (at time zero) and offset (28.5 seconds) for high (red) vs. low (blue) current. C, (upper) Mean firing rates for all single units in each area as a function of time in the trial during high current stimulation (blue bars: pre-stim baseline, orange bars: electrical stimulation, yellow bars: 0-30 seconds after electrical stimulation offset, purple bars: 30-60 seconds after electrical stimulation offset, green bars: 60-90 seconds after electrical stimulation offset). (lower) same as upper subplot, but for low current stimulation. Mean and 99^th^ confidence interval. D, Mean dB change in power as a function of time since electrical stimulation for all high current trials in area 8A. F, Same as D, but for low current. E, Same as D, but highlighting spectral modulation during the period of electrical stimulation (0-28.5 seconds). Mean +/− 1SEM. G, same as D, but highlighting spectral modulation during the post-1 period of electrical stimulation (0-30 seconds post electrical stimulation offset). Mean +/− 1SEM.

LFP power was modulated by electrical stimulation (shown for area 8A in Figure 9 D and F and for the other areas in Supplemental Figure 4). High-current, but not low-current, stimulation significantly reduced slow frequency power for up to 85 seconds post electrical stimulation onset relative to pre stimulation baseline (Figure 9D and Supplemental Figure 4, P<0.01, cluster-based non-parametric randomization test). High-current stimulation also significantly increased higher frequency power during and after stimulation (Figure 9D and Supplemental Figure 4, P<0.01, cluster-based non-parametric randomization test). During stimulation, the enhancement had a well-defined peak in the alpha/beta band in area 8A and PFC (vertical dotted lines in Figure 9E indicate peaks at 13 Hz in 8A and 15 Hz in PFC). They were absent in the 0-30 second post-stimulation offset interval. The reduction in slow frequency power persisted for 85 seconds post-stimulation (Figure 9G). All of the effects were weak or absent for low current stimulation (Figure 9F).

## Discussion

Our study provides new details on the neurophysiological effects of propofol-induced unconsciousness. Just prior to unconsciousness, there was an increase in alpha/beta power in frontal cortex. By contrast, alpha/beta power in posterior cortex transiently increased and then decreased relative to the awake state. During unconsciousness, there was a widespread decrease in high-frequency (greater than 50 Hz) power and an increase in slow-frequency (~1 Hz and lower) power. Spike rates decreased and spiking became coupled to the slow-frequency oscillations. There was an increase in slow-frequency cortico-cortical and thalamocortical phase synchronization. In frontal cortex alone, there was an increase alpha/beta (~8-30 Hz) power and cortico-cortical and corticothlamic synchronization. These effects were distributed differently across cortical layers. In superficial layers (2/3), there was a stronger suppression of gamma and spiking during unconsciousness. Deep layers (5/6) showed a stronger increase in slow-frequency power.

Our results agree with results in humans that have related slow frequency oscillations with propofol-induced unconsciousness. We also observed a profound decrease in cortical spiking and an increase of its phase-locking with slow-frequency oscillations. The evidence that this fragmented cortex or disconnected the frontal and parietal cortex is less well-supported. This may cause unconsciousness by disrupting cortical communication, especially in the theta, beta, and gamma ranges associated with cognition and consciousness (Bastos et al., 2015; Brown et al., 2010; Buschman and Miller, 2007; Cimenser et al., 2011c; Fiebelkorn and Kastner, 2019; Lakatos et al., 2008; Lewis et al., 2012a; Miller et al., 2018; Vijayan et al., 2013). We also found different effects across layers that support other theories. Dehaene and Changeux (2011) proposed that the conscious state relies on “broadcasting” of cortical activity throughout cortex via long-range superficial connections. Aru, Larkum, and others (2019) suggested that consciousness relies on cortico-thalamic broadcasting which depends on integrity of sub-cortical projection neurons in deep layers.

The thalamus is known to contribute to cortical dynamics in the awake state (Fiebelkorn and Kastner, 2019; Fiebelkorn et al., 2019). The ILN of the central thalamic nucleus has diffuse cortical connections posited to mediate the global binding needed for consciousness (Llinás et al., 1998). Thalamic stimulation has improved behavioral performance of a minimally conscious patient (Schiff et al., 2007). Correspondingly, we found that stimulation of the central thalamus caused monkeys to regain arousal, similar to a recent report (Redinbaugh et al., 2020). The stimulation increased cortical spike rates, diminished slow-frequency power, and re-instated higher-frequency power. Our studies are consistent with rodent studies that have shown recovery of consciousness and awake-like cortical dynamics after stimulation in a variety of brain regions (Pillay et al., 2014; Solt et al., 2014).

Our results are generally in agreement with a recent study that used similar techniques in NHPs (Redinbaugh et al., 2020). Redinbaugh et al (2020) reported that the unconscious state was not characterized by increased slow-frequency synchronization between cortex and thalamus. We found that increased slow-frequency power and coherence is an important feature of the propofol-induced unconscious state in all areas. Redinbaugh et al (2020) only found decreases in spike rate in deep cortical layers. We found a stronger decrease in superficial layers. These are the first studies of this kind. Further experimentation can resolve these details. Yet, there were key differences between our studies. Redinbaugh et al. (2020) administered ketamine before they began experimentation began with propofol. In contrast, our anesthetic regime consisted of only propofol administration. They only compared awake to a stable state of unconsciousness and this could not study the transitions into and out of unconsciousness as we did.

Thus, propofol likely renders unconsciousness by disrupting intracortical and thalamocortical communication through decrease spiking and enhanced slow-frequency power/synchrony. This leads to low-spiking Down-states and loss of the higher frequency coherence thought to integrate cortical information (Baars et al., 2013; Cimenser et al., 2011c; Crick and Koch, 1990; Dehaene and Changeux, 2011; Dehaene and Naccache, 2001; Demertzi et al., 2019; Engel et al., 1999; Lee et al., 2013; Purdon et al., 2013). The slow-frequency oscillations could be due to direct inhibition of brainstem, thalamus and cortex (Brown et al 2010; Brown et al 2011; Brown et al. 2018).

---

**Supplemental Figure 1.**
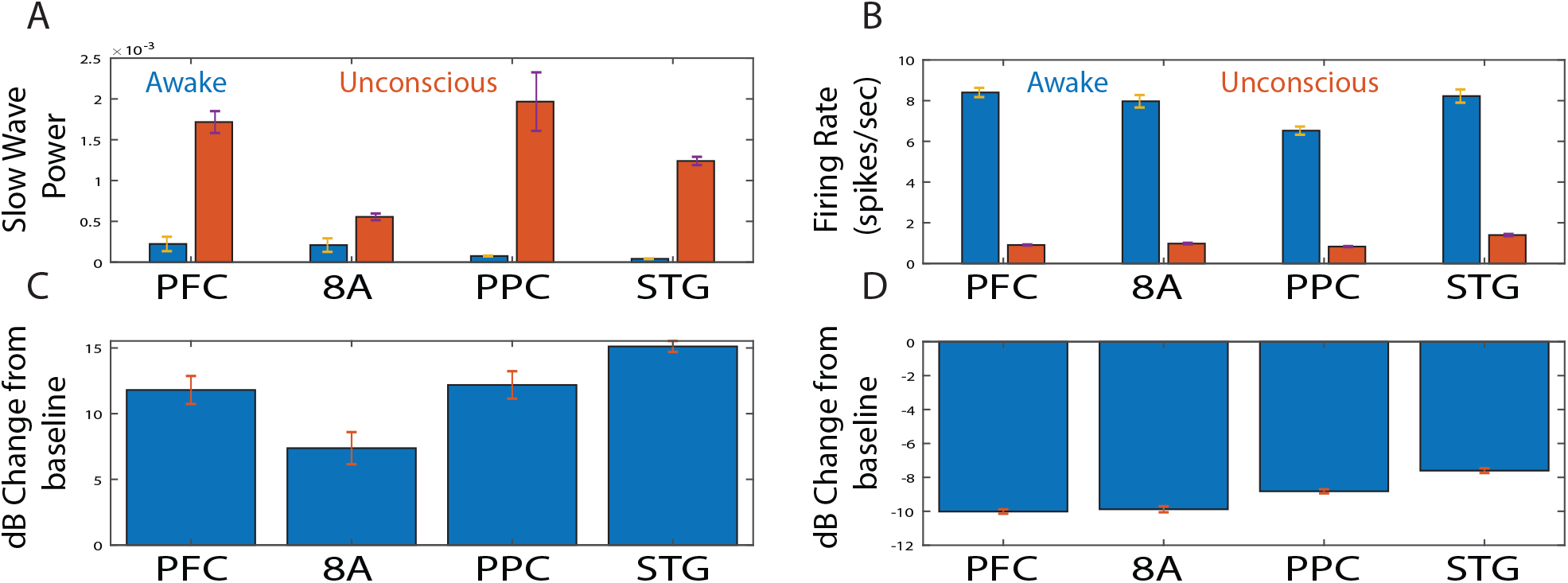
Effect sizes for change in power vs. spiking during unconsciousness. A, Slow wave (0.1-2 Hz) power during the awake (blue bars) vs. unconscious (orange bars) states. Mean +/− 1SEM across sessions. B, Firing rates during the awake (blue bars) vs. unconscious (orange bars) states. Mean +/− 1SEM across all units per area. B, The dB change in SF power, averaged across all sessions. Mean +/− 1SEM across sessions. D, The dB change in spiking, average across all units per area. Mean +/− 1SEM across all units per area.

**Supplemental Figure 2.**
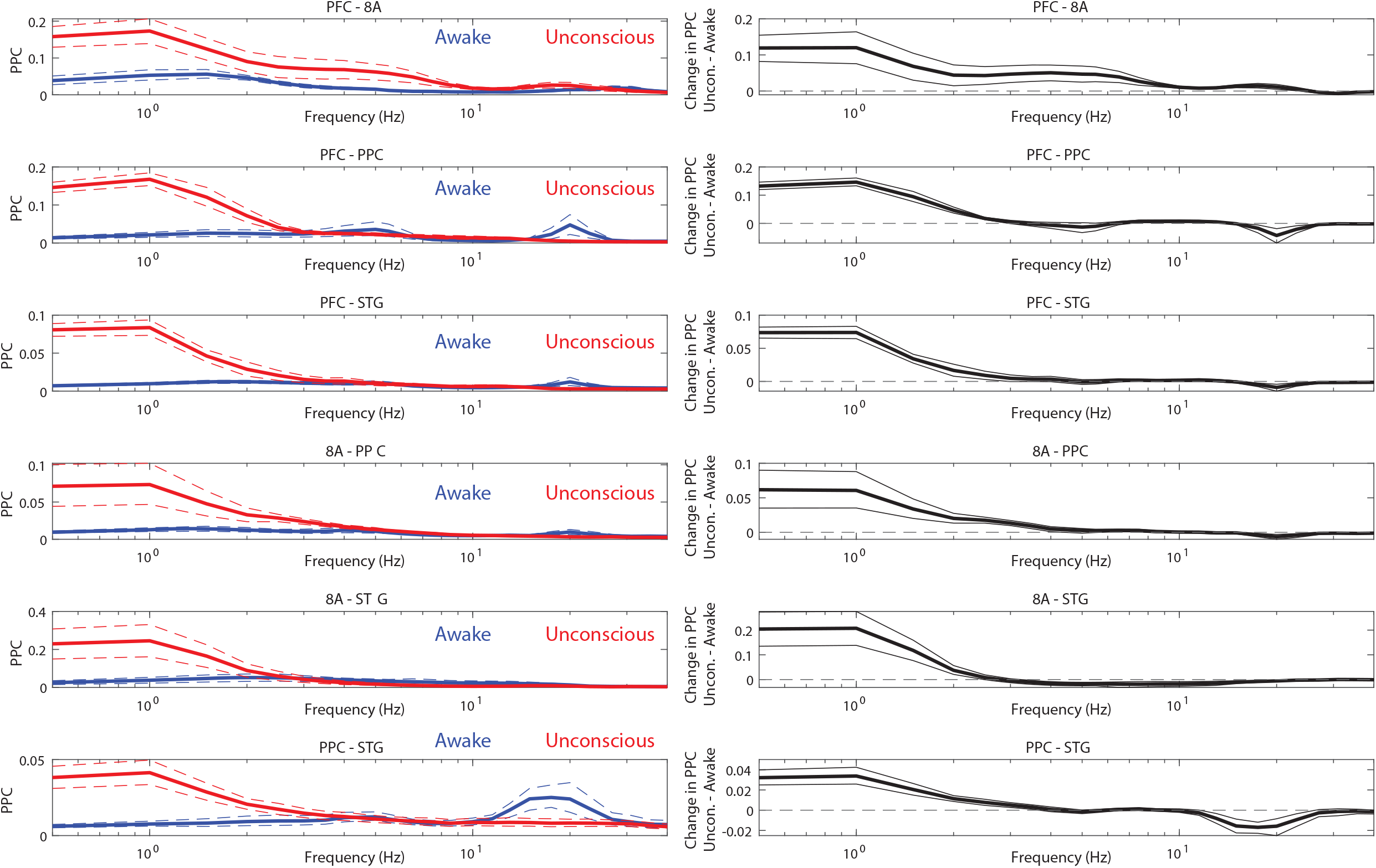
Effect sizes for change in cortico-cortical coherence during awake vs. unconsciousness. (left panels) Average Pairwise Phase Consistency (PPC) during the pre-propofol awake (blue) vs. unconscious (red) states. (right subpanels). Average PPC difference (unconscious – conscious). Mean is displayed in bold lines and the 99^th^ confidence interval of the mean is shown in dotted lines.

**Supplemental Figure 3.**
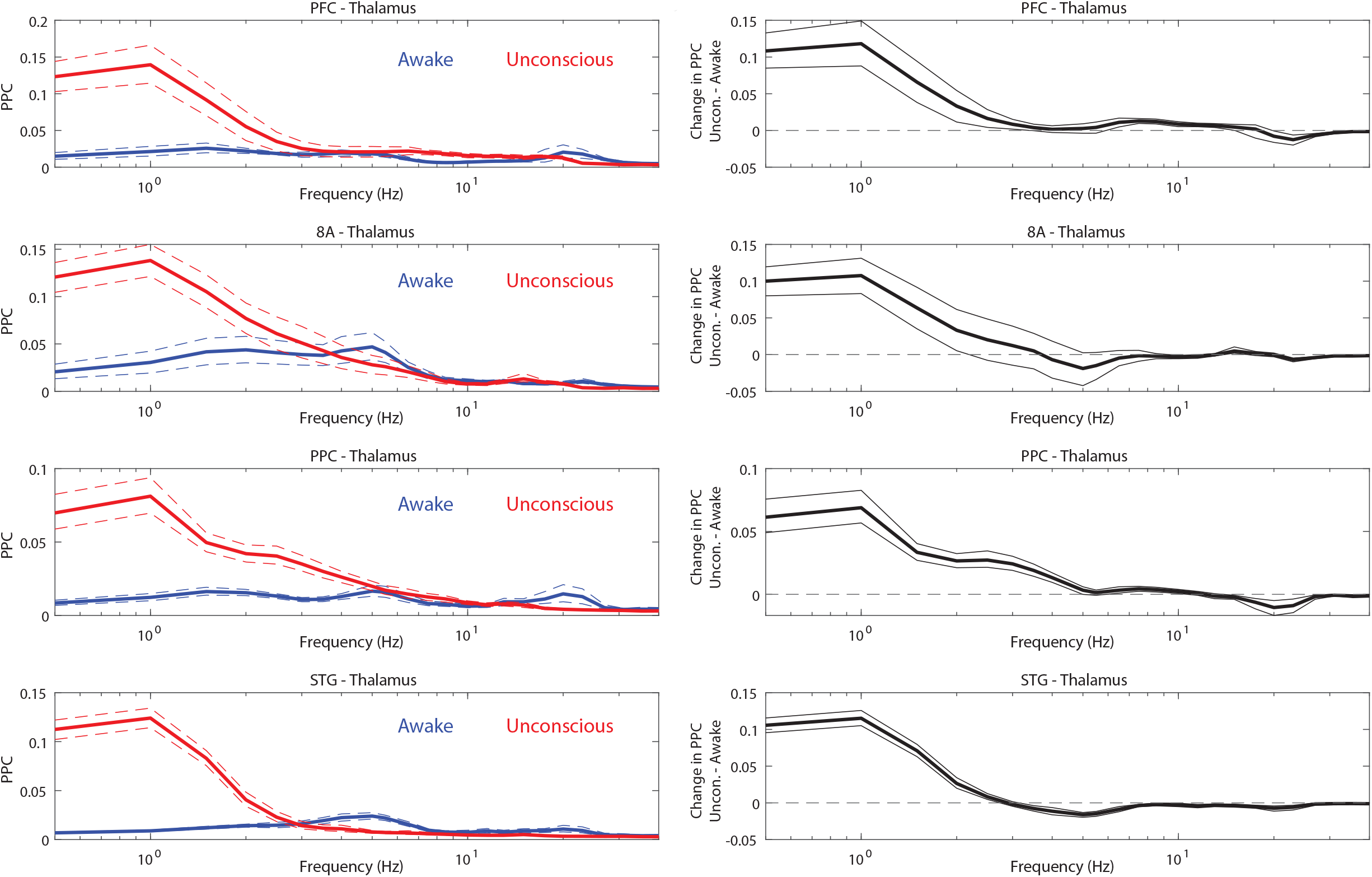
Effect sizes for change in thalamos-cortical coherence during awake vs. unconsciousness. Same as Supplemental Figure 2 but for thalamocortical interactions.

**Supplemental Figure 4.**
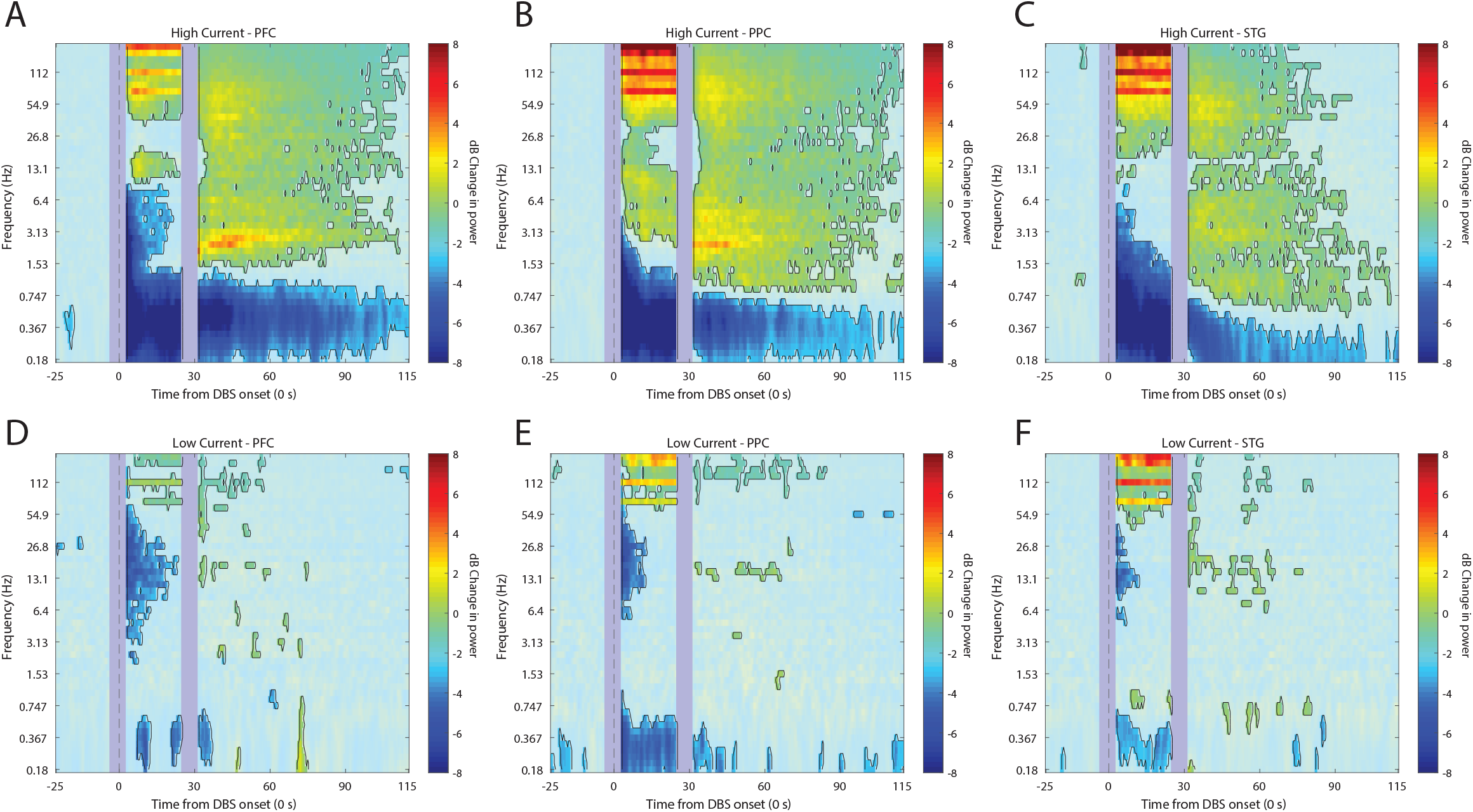
Effects of thalamic electrical stimulation on cortical power. A/B/C Mean dB change in power as a function of time since electrical stimulation for all high current trials in areas PFC, PPC, and STG, respectively. D/E/F, Same as A/B/C, but for low current. Significant increases or decreases from pre-stimulation baseline are marked with opaque colors and are highlighted.

## Materials and Methods

### Experimental Subjects and vascular access port

Two rhesus macaques (*Macaca mulatta*) aged 14 years (MJ, male, ~13.0 kg) and 8 years (Monkey LM, female, ~6.6 kg) participated in these experiments. Both animals were pair-housed on 12-hr day/night cycles and maintained in a temperature-controlled environment (80°F). Each monkey was surgically implanted with a subcutaneous vascular access port (Model CP-6, Norfolk Access Technologies, Skokie, IL) at the cervicothoracic junction of the neck with the catheter tip reaching the termination of the superior vena cava via the external jugular vein.

### Neural recordings in cortex

Neurophysiology For recordings in cortex, monkeys were chronically implanted with four 8×8 iridium-oxide contact microelectrode arrays (“Utah arrays”, MultiPort: 1.0 mm shank length, 400 μm spacing, Blackrock Microsystems, Salt Lake City, UT), for a total of 256 electrodes. Arrays were implanted in the prefrontal (area 46 ventral and 8A), posterior parietal (area 7A/B), and temporal-auditory (caudal parabelt area STG, Superior Temporal Gyrus) cortices. Specific anatomical targeting utilized structural MRIs of each animal and a macaque reference atlas, as well as visualization of key landmarks on surgical implantation (McLaren et al., 2009). For Utah array recordings, area 8A and PFC were ground and referenced to a common subdural site. Area STG and PPC also shared a common ground/reference channel which was also subdural. LFPs were recorded at 30 kHz and filtered online via a lowpass 250Hz software filter and downsampled to 1 kHz. Spiking activity was recorded by sampling the raw analog signal at 30 kHz, bandpass filtered from 250Hz-5kHz, and manually thresholding. Blackrock Cereplex E headstages were utilized for digital recording via 2–3 synchronized Blackrock Cerebus Digital Acquisition systems. Single units were sorted manually offline using principal component analysis with commercially available software (Offline Sorter v4, Plexon Inc., Dallas, TX). All other pre-processing and analyses were performed with Matlab (The Mathworks, Inc, Natick, MA).

To ensure signals recorded on the multiple data acquisition systems remained synchronized with zero offset, a synchronization test signal with locally unique temporal structure was recorded simultaneously on one auxiliary analog channel of each system. The relative timing of this test signal between each system’s recorded datafile was measured offline at regular intervals throughout the entire recording session. Any measured timing offsets between datafiles were corrected by appropriately shifting spike and event code timestamps in time, and by linearly interpolating analog signals to a common time base.

### Neural recordings and electrode targeting in thalamus

After 10-20 sessions of cortex only recordings, monkeys were chronically implanted with four 6-8 channel recording/stimulating electrodes (0.5 mm contacts with 0.5 mm intercontact-spacing, NuMed Inc, Hopkington, NY) bilaterally targeting the intralaminar nuclei of the central thalamus. Thalamic recordings were referenced to the monkey’s titanium headpost. In offline analysis, thalamic sites were re-referenced to a bipolar montage prior to further analysis. LFPs were recorded via a separate analog front end amplifier and an additional identical digital acquisition system, synchronized to the digital acquisition systems utilized for cortical recordings. LFPs were similarly recorded at 30 kHz and filtered online via a lowpass 250Hz software filter and downsampled to 1 kHz.

A specialized anatomical localization and insertion protocol utilizing serial intraoperative MRIs was developed in order to allow for precise subcortical targeting of the electrodes along the long axis of the central lateral nucleus and extending ventrally to the centromedian and parafasicular nuclei. Custom-made carbon PEEK recording chambers were affixed to the skull with acrylic and ceramic screws stereotaxically determined to target the central thalamus. Recording chamber grids with 1mm grid holes were inserted into the chambers, filled with sterile saline, and the monkeys were imaged by 3T MRI. After confirmation of the appropriate grid holes targeting the thalamic structures of interest, monkeys were generally anesthetized and brought to the operating facility where a small-bore craniotomy (<2mm) was performed at the appropriate grid holes. The grid was replaced in the chamber and the monkeys were transferred to the imaging facility under general anesthesia. They were then administered a gadofosveset trisodium contrast agent to highlight vasculature obstructing the trajectory to thalamus (e.g. thalamostriate vein). In the MRI suite, a stylette cannulae was inserted into the relevant grid holes and lowered several mm into cortex and one set of 0.5 mm resolution images was obtained. Upon confirmation of correct trajectory on MRI, the stylette-cannulae were lowered to their final position, with the tip approximating the thalamus. The stylettes were then removed, and electrodes of marked length lowered to the depth of the cannulae. Following another MRI-based measurement (scan 2) with the electrodes still in the cannulae, the electrodes were lowered to their final positions within the thalamus and reimaged (scan 3). Upon final assessment of correct localization, the probes were fixed in place and the chamber sealed with acrylic. Histological staining with acetylcholinesterase was used to confirm exact electrode contact locations within and outside the central thalamus of both monkeys.

### Experimental Procedures

Considering both the “propofol cortex sessions” and the “thalamus wakeup sessions” (see below for details), a total of 43 sessions were analyzed (N=22 from monkey 1 and N = 21 from monkey 2).

On a given experimental session, monkeys were head-fixed via a titanium headpost and placed in noise isolation chambers with masking white noise (50 dB). We ran two sets of experimental sessions. The first set of sessions consisted of neurophysiological recordings from cortex only. We refer to these as “propofol cortex sessions”. A total of 21 sessions (N=11 from monkey 1, N = 10 from monkey 2) were used. These sessions proceeded as follows: first, a period of 15 - 90 minutes of awake baseline electrophysiological recordings were recorded. Next, propofol was intravenously infused via a computer-controlled syringe pump (PHD ULTRA 4400, Harvard Apparatus, Holliston, MA). The infusion protocol was stepped such that unconsciousness was induced via a higher rate infusion (285 μg/kg/min for monkey MJ; 580 μg/kg/min for monkey LM) for 20 minutes before dropping to a GA maintenance dose (142.5 μg/kg/min for monkey MJ; 320 μg/kg/min for monkey LM) for an additional 40 minutes.

The second set of experimental sessions occurred after we implanted the thalamic recording/stimulation electrodes (<u>d</u>eep <u>b</u>rain <u>s</u>timulating electrodes, or DBS). We refer to these as the “thalamus wakeup sessions”. After the initial 20 minutes of propofol infusion, DBS stimulation trials began. A total of 22 sessions (N=11 from monkey 1, N = 11 from monkey 2) were analyzed with thalamic stimulation.

Heart rate and oxygen saturation were monitored continuously and recorded throughout all phases of experiments using clinical-grade pulse oximetry (Model 7500, Nonin Medical, Inc., Plymouth, MN). SpO_2_ values were maintained at values above 93% for each of the recording sessions.

Infrared monitoring tracked facial movements and pupil size (Eyelink 1000 Plus, SR-Research, Ontario, CA) throughout the course of the experiments. Loss of consciousness (LOC) was deemed by the timestamp of the moment of eyes-closing that persisted for the remainder of the infusion. Recovery of consciousness (ROC) was classified as the timestamp of the first to occur between eyes reopening or regaining of motor activity following drug infusion cessation. Animals regained consciousness after the maintenance infusion was terminated and were monitored for an additional period before being returned to their home cage. To ensure propofol clearance from tissues and physiological recovery, experiments were never repeated on subsequent days. All procedures followed the guidelines of the MIT Animal Care and Use Committee and the US National Institutes of Health.

### Thalamic stimulation procedure

For electrical stimulation of thalamic electrodes, we adapted electrical stimulation parameters previously shown to cause behavioral improvements in coma patients and awake, behaving monkeys (Baker et al., 2016; Schiff et al., 2007b). We unilaterally delivered 180 Hz bipolar, biphasic, square wave pulses (0.5-2.5 milliAmps) between 6-8 contacts on monkey MJ and 6 contacts on monkey LM. Five minutes into the maintenance anesthesia dose (20 minutes from infusion start), 30-second “trials” of electrical stimulation were delivered as the propofol infusion continued. DBS trials were separated by 2 minutes intervals, except the 4th and 5th stimulation runs, which were separated by a 5-minute inter-trial interval. These DBS washout periods sufficiently allowed for reestablishment of the behaviorally-judged anesthetized state (e.g. loss of puff responses). We delivered between 0.5-2.5 mAmp of current. In early pilot experiments, different currents, frequencies, waveform shapes, and electrode combinations were screened for eliciting arousal. We chose a set of parameters that was effective at eliciting arousal, but minimal in current and number of stimulated electrodes, for the final experiments reported here.

### Wake up Score

One of the authors (MM) who was present during all DBS sessions performed a behavioral score for each DBS trial for the degree to which the electrical stimulation induced changes in arousal. This numerical “wake up score” is loosely inspired by the Glasgow coma scale. Like the Glasgow coma scale, it separately scores each of the behavioral components, then sums them into a single overall score for each DBS trial. The components are: (1) Spontaneous eye opening (0–2): 0 = eyes closed, 1 = one or both eyes slightly open, 2 = one or both eyes fully open; (2) Responses to external stimuli (airpuffs directed at eye/face; 0–2): 0 = no response, 1 = occasional blinking not necessarily in response to puffs, 2 = clear response to airpuffs; (3) Face/body movements (0–2): 0 = none, 1 = some mouth/jaw movement, 2 = arm/full-body movement. The final waking score for each trial is simply the sum of these three components. Note that the actual Glasgow scale combines components 1 & 2 into a single “eye opening” score, but empirically in this data, spontaneous eye opening and blink responses to airpuffs seem to occur somewhat independently rather than on a single continuum.

### Histology

At the end of the recording sessions, the animals were euthanized for histological confirmation of thalamic sites, as previously described (Wu and Kaas, 1999). Briefly, monkeys were given a lethal dose of sodium pentobarbital. When they became areflexive, they were perfused transcardially with PBS, followed first by a cold solution of 4% paraformaldehyde and next by a mixed solution of 4% paraformaldehyde and 10% sucrose. Blocks of brain and spinal cord were removed and stored overnight in 30% sucrose at 5°C before cutting. Sections were processed for acetylcholinesterase. Anatomical localization of electrodes was determined by histological examination of brain tissue. It was not necessary to create electrolytic lesions prior to histology, because the thalamic electrodes were wide enough (0.5mm diameter) to be unambiguously identified in the anatomical sections.

### Data Preprocessing, general

Single units were sorted manually offline using principal component analysis with commercially available software (Offline Sorter v4, Plexon Inc., Dallas, TX). All other pre-processing and analyses were performed with Matlab (The Mathworks, Inc, Natick, MA).

### Data Preprocessing, electrical stimulation LFP

Electrical stimulation generally produced artifacts that were highly correlated across channels in a stereotyped manner. We removed these using zero-phase component analysis (ZCA) whitening (Eldar and Oppenheim, 2003). ZCA whitening is the linear whitening transformation that minimizes the mean squared error between the original and whitened signals. Intuitively, it estimates the across-channel correlations induced by DBS stimulation, and removes them from the data. First, we extracted the spiking band from the raw 30 kHz analog signals by band-pass filtering at 300–4500 Hz with a zero-phase fourth-order Butterworth filter. Next, we estimated the cross-channel covariance matrix Σ from the filtered signals during the DBS stimulation periods (from a subset of 100,000 randomly sampled time points, for computational efficiency). From the estimated covariance, we computed the “whitening matrix”: W=Σ^(−1/2). We then normalized each column of W by its diagonal (variance) value, so that the resulting matrix W^* will remove DBS-induced correlations, but not change the amplitude of individual channels. We removed the DBS stimulation artifacts by multiplying the full matrix of filtered signals by the modified whitening matrix: 〖D_denoised=D_raw W〗^*. Finally, we computed the mean and standard deviation of each denoised channel, thresholded each at −4.5 SD, and extracted spike timestamps and waveforms around each threshold crossing. Extracted spikes were sorted into units using principal component analysis in commercially-available software (Offline Sorter v4, Plexon Inc., Dallas, TX). Only spikes whose average waveform during electrical stimulation matched the waveform outside periods of electrical stimulation (Pearson correlation coefficient greater than or equal to 0.99) were included for analysis.

### Methods, LFP spectral analysis and statistics

Local Field Potentials (LFPs) were analyzed using the Fieldtrip toolbox (http://www.fieldtriptoolbox.org/) (Oostenveld et al., 2011). We tested whether specific oscillations in different areas relative to drug onset (or electrical stimulation onset) were modulated in power. For each channel on each array or thalamic probe, we computed a time-frequency decomposition. For propofol only sessions, we calculated power in sliding windows of duration 10 seconds with a hanning taper, to deliver 0.1 Hz Spectra resolution. We calculated power across logarithmically spaced frequencies 0.178-200 Hz (Figure 2).

We calculated the change in power between drug and pre-drug baseline in decibel (dB) units. In other words, we applied the following transformation to the raw power values:

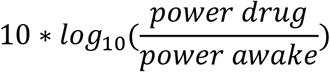

We took the pre-drug baseline to be the 2 minutes immediately before drug onset, and the drug period to be every second after drug onset.

For cortico-cortical coherence analysis, we first re-referenced data to a local bipolar montage at 1600um distance. We then calculated coherence between inter-areal bipolar sites using the multitaper method with 1 Hz spectral smoothing to estimate power in 0.5 Hz intervals from 1 to 200 Hz using 5 second windows with 75% overlap. For thalamocortical coherence analysis, we re-referenced cortical recordings as above, and we re-referenced thalamus data to a local bipolar montage at 1500um distance. We calculated power from 60 seconds prior to propofol onset to 1200 seconds post onset (20 minutes, when the first DBS stimulation trials began). Each window of analysis for coherence consisted of 50 seconds of data, which were further subdivided into 2 second windows to create enough pseudo-trial repetitions to estimate coherence.

For cortex power modulation with thalamic electrical stimulation, we calculated power using a sliding window approach time locked to the onset of electrical stimulation. Power was calculated from 25 seconds pre-stimulation to 150 seconds post-stimulation with 0.5 second intervals, and with 5 second analysis windows. We calculated change in power relative to pre-stimulation baseline, which was the average power from 25 seconds to 3 seconds prior to stimulation.

### Statistics

We determined whether there were differences from baseline in power and coherence by using a non-parametric cluster-based randomization test (Oostenveld et al., 2011). For each session, we realized the null hypothesis that power in the baseline and power in the drug period were the same. To this end, we randomly exchanged baseline-transformed time-frequency power estimates between the baseline and drug windows. We extracted the largest cluster (continuous tiles in time-frequency space) to pass a first level significance threshold, by applying a t-test and thresholding all significant bins p<0.01, uncorrected. We performed this randomization 1,000 times. The empirically observed clusters were compared to this randomization distribution to assess significance at p=0.01, adjusted for multiple comparisons across sessions.

To calculate the effect of electrical thalamic stimulation on cortical power, we applied a similar transformation, only this time the baseline was the 30 seconds of data immediately preceding each trial of stimulation:

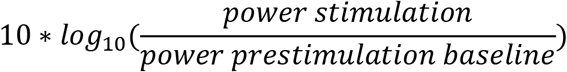

We then repeated the same test outlined above, only now randomizing bins before/during/after stimulation onset to create the null hypothesis. There were stimulation onset and offset artifacts. We removed the times around onset/offset +/− 3 seconds prior to performing this randomization test. As a result, these artifact times are omitted from the analysis and figure.

### Methods, state-space modeling of physiological signals

To characterize physiologically the transition from consciousness to unconsciousness, we measured heart rates, muscle tone with electromyography (EMG), and blood oxygenation. These signals were pre-processed in the following manner. Heart rates and SPO2 signals were averaged using non-overlapping windows of one second. The EMG signals were z-scored by subtracting its mean and dividing by its session-wide standard deviation. Its variance was computed using one second, non-overlapping windows. From EMG signals measured from two electrodes adjacent to the eyes, we extracted evoked puff responses (eyeblinks) using the following procedure. We computed the average of the EMG signal within a 50-150 millisecond time window, following an air-puff stimulus, for each individual session. We subtracted the mean, and divided this time-series by its session-wide standard deviation. Then, we computed a moving average over a 30 second window to obtain a continuous estimate of this response.

To quantify the change in these signals after the propofol infusion, we used a state space model approach. We assume that the log of the EMG variance, the blood oxygenation levels, and the heart rate signals can be modeled under the linear gaussian state-space model framework. Since there may be missing observations, we use the following formulation:

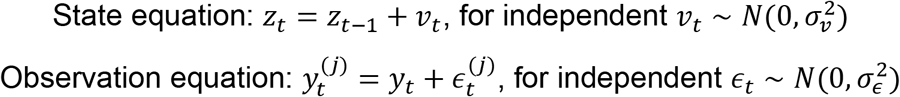

Under this formulation, we use all physiological measurements across sessions, for each type of physiological measurement. There are J maximum number of observations at each timepoint, each observation has its own equation, and all observations share the same state. This allows for estimating the state of the latent process while having missing observations in the data. Since *z*_*t*_, 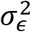 and 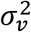 are unknown, we estimate these parameters through maximum-likelihood via the Expectation-Maximization (EM) algorithm (Dempster, 1977). A derivation of our EM algorithm is included in Appendix A.

To compare these physiological measurements across different times, we computed the probability that a measurement at time *t* was lower than measurements at all previous time points. We performed this comparison since these measurements seem to decrease after the first propofol infusion. We computed this probability for all time-points using a Monte Carlo algorithm detailed in (Smith et al., 2005) We considered a result to be statistically significant if the posterior probability was greater than 0.99.

### Testing whether spikes become coupled to the LFP slow oscillation phase

We used a generalized linear model (GLM) framework to test whether spike trains from individual neurons could be modeled as a linear combination of the slow-frequency/delta LFP phase.

Spike rates were modeled as

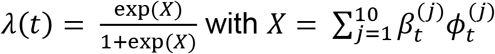

Where *λ*(*t*) is the instantaneous spike rate of each individual neuron at time t, 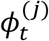 is an indicator random variable that takes value 1 if the phase of the LFP oscillation at time *t* is in phase bin *j,* and 0 otherwise. This defines a GLM with a binomial likelihood and a logit link function (McCullagh and Nelder, 1989; Truccolo et al., 2005). For GLM fitting, we used custom software for performing regression with Truncated Regularized Iteratively Reweighted Least Squares (Komarek, 2005). Goodness-of-fit was assessed using Kolmogorov-Smirnov (K-S) tests based on the time-rescaling theorem (Brown et al., 2002)

Phase bins were computed using the following procedure. First, we band-passed the LFP in the slow-frequency/delta band (0.3-3Hz). We then computed the Hilbert transform to extract a measure of the continuous analytic phase. Finally, we binned these phase estimates into ten linearly spaced bins ranging from −*π* to *π*. A one-hot encoding of these phase bins was used as the design matrix for the model.

### Testing whether there is an increase in phase-modulation after loss of consciousness

To test whether there was an increase in phase-modulation after LOC, we fit GLMs to 20 minutes of data during the pre-drug awake state, and to 20 minutes of data after LOC. We defined phase-modulated neurons as neurons whose GLMs had a good fit to the data. We then compared the proportion of phase-modulated neurons post-LOC to the proportion of phase-modulated neurons during the pre-drug awake state by performing a Bayesian comparison using the beta-binomial model (DeGroot et al., 2002; Solt et al., 2011). We assumed a binomial model as the likelihood function for the proportion of phase-modulated neurons for each condition. We used a uniform prior on the interval (0,1) as the prior density, and a beta posterior density due to conjugacy. We then computed the posterior density for the difference in the proportion of phase-modulated neurons (post-LOC minus awake). Using a Monte Carlo procedure and 10,000 Monte Carlo samples, we computed the probability that the proportion of phase-modulated neurons post-LOC was greater than the proportion of phase-modulated neurons during the pre-drug awake state, as in Chemali et al., 2012. We considered a result to be statistically significant based on the 95% Bayesian credibility intervals for comparing these two conditions if zero was not in the interval. We considered a result to be statistically significant based on the posterior probability if this value was greater than 0.95.

## Acknowledgements

We would like to thank Simon Kornblith for help with data analysis. These studies were supported by NIMH R01MH11559, NIGM P01GM118269 and The MIT Picower Institute Innovation Fund.

